# Thymically-derived Foxp3+ regulatory T cells are the primary regulators of type 1 diabetes in the non-obese diabetic mouse model

**DOI:** 10.1101/644229

**Authors:** Daniel R. Holohan, Frédéric Van Gool, Jeffrey A. Bluestone

## Abstract

Regulatory T cells (Tregs) are an immunosuppressive population that are identified based on the stable expression of the fate-determining transcription factor forkhead box P3 (Foxp3). Tregs can be divided into distinct subsets based on whether they developed in the thymus (tTregs) or in the periphery (pTregs). Whether there are unique functional roles that distinguish pTregs and tTregs remains largely unclear. To elucidate these functions, efforts have been made to specifically identify and modify individual Treg subsets. Deletion of the conserved non-coding sequence (CNS)1 in the *Foxp3* locus leads to selective impairment of pTreg generation without disrupting tTreg generation in the C57BL/6J background. Using CRISPR-Cas9 genome editing technology, we removed the *Foxp3* CNS1 region in the non-obese diabetic (NOD) mouse model of spontaneous type 1 diabetes mellitus (T1D) to determine if pTregs contribute to autoimmune regulation. Deletion of CNS1 impaired *in vitro* induction of Foxp3 in naïve NOD CD4+ T cells, but it did not alter Tregs in most lymphoid and non-lymphoid tissues analyzed except for the large intestine lamina propria, where a small but significant decrease in RORγt+ Tregs and corresponding increase in Helios+ Tregs was observed in NOD CNS1^−/−^ mice. CNS1 deletion also did not alter the development of T1D or glucose tolerance despite increased pancreatic insulitis in pre-diabetic female NOD CNS1^−/−^ mice. CNS1 Furthermore, the proportions of autoreactive Tregs and conventional T cells (Tconvs) within pancreatic islets were unchanged. These results suggest that pTregs dependent on the *Foxp3* CNS1 region are not the dominant regulatory population controlling T1D in the NOD mouse model.

## Introduction

Regulatory T cells (Tregs) are an immunosuppressive subset of CD4+ T cells important for the maintenance of self-tolerance [1,2]. Tregs were initially marked by their high expression of the α chain (CD25) of the interleukin-2 (IL-2) receptor and then later identified as constitutively expressing the fate-determining forkhead box P3 (Foxp3) transcription factor, which confers stable cell identity and suppressive function [3–5]. Mutations in Foxp3 or Treg deletion in adult mice leads to systemic autoimmune pathology in human and mice, including type 1 diabetes (T1D) [6–10].

A number of studies have shown that Tregs can develop both in the thymus (tTreg) and post-thymically in the periphery (pTreg), especially in the gut. During negative selection in the thymus, high affinity, self-reactive cells can escape clonal deletion by differentiating into Tregs, which is facilitated through CD28 costimulation and Foxp3 expression [11,12]. In contrast, naïve conventional T cells (Tconvs) exposed to self-antigens and the microbiome in the periphery can differentiate into Tregs, especially in mucosal tissues and in context of transforming growth factor-beta (TGF-β) and other pro-Treg environmental factors [13–16]. Direct evidence of pTreg generation can be seen when antigens were administered in the periphery by osmotic pumps [17], oral administration [18], or targeted antigen delivery to dendritic cells [19].

Previous studies have shown that Tregs are defective in both mice and humans that develop T1D. Thymic development and the Treg repertoire have been shown to be altered in NOD mice [20,21], and reducing T-cell receptor (TCR) diversity in NOD mice alleviates T1D, connected to a lack of insulin beta chain (InsB:9-23) autoreactive T cells [22]. In fact, recent studies have identified islet-antigen-specific Tregs that can control disease progression [23]. However, it remains unclear where these Tregs are generated and which subset of Tregs are defective. Finally, fusion peptides made by post-translational modifications of insulin derivatives and other proteins also create neoantigens that more strongly bind to potentially autoreactive cells in NOD mice [24–26] and humans [25,27–29]. Although the role of these neoantigens and their recognition by Tconvs and Tregs remain unclear, their existence suggests that peripherally-derived antigens not typically found in the thymus might drive T1D and potentially pTreg generation. Interestingly, adoptive transfer experiments have shown non-redundant roles for tTregs and pTregs in maintaining tolerance [30].

Identification of phenotypic or molecular markers that distinguish tTregs from pTregs *in vivo* has proven more challenging. Expression of the surface marker Neuropilin-1 (Nrp1, also known as CD304) [31,32] and the transcription factor Helios [33] have been linked to tTregs; however, Helios and Nrp1 can also be markers of activation [34–36], and Nrp1 expression has been observed in some pTregs generated during adoptive transfer experiments and in tissue resident Tregs [37]. As a result, efforts have been made to identify distinct regulatory elements that differentially control Foxp3 expression in the different Treg subsets. Genetic studies by Rudensky and colleagues showed that there are 3 conserved non-coding sequence (CNS) elements, termed CNS1, CNS2, and CNS3, upstream of the *Foxp3* promoter that control Foxp3 expression. One element, CNS1, was shown in C57BL/6J (B6) mice to selectively affect pTreg development without perturbing thymic expression of Foxp3 [38]. This region contains Smad-binding motifs downstream of the TGF-β signaling pathway, which has been implicated in the development of induced Tregs (iTregs) differentiated from naïve CD4+ T cells *in vitro* and gut-derived pTregs through mechanisms such as the activation of latent TGF-β through interactions with integrin α_v_β_8_[39]. Administration of TGF-β *in vivo* can also enhance the antigen-specific generation of pTregs [19,40]. B6 CNS1^−/−^ blocked increases in Treg percentages in peripheral tissues with age, resulting in Th2-mediated pathology in the lungs and intestinal tract [41]. Continued characterization of the B6 CNS1^−/−^ mouse revealed defects in pTreg generation and gut microbiome colonization in CNS1^−/−^ mice [42,43] as well as in maternal-fetal tolerance [44].

There have been limited studies on the relative roles of pTregs and tTregs in controlling autoimmunity. Studies of experimental allergic encephalomyelitis (EAE) in a B6 CNS1^−/−^ mouse model suggested that pTreg deficiency did not exacerbate disease [41], and adoptive transfer of TCR transgenic tTregs but not pTregs protected mice from EAE [31]. These results are consistent with another group who showed that a Smad3 binding mutation within *Foxp3* CNS1 of NOD mice did not affect susceptibility to colitis, and although not stated explicitly, the authors did not note an increased incidence of T1D [45]. In contrast, other studies have shown that in NOD CD28^−/−^ mice, adoptive transfer of islet antigen-specific pTregs was equally effective as tTregs in controlling T1D (31), and in one study, co-transfer of TCR transgenic BDC2.5tg+ pTregs with BDC2.5tg+ CD4+ T was sufficient to control T1D in NOD RAG1^−/−^ mice [46]. In sum, these results suggest that tTregs are most critical in preventing and reversing autoimmunity and that in most cases pTregs are not effective. However, in the case of unmanipulated, polyclonal Tregs, the results are less clear.

Therefore, the goal of this study was to address the role of polyclonal pTregs in controlling autoimmune diabetes. pTreg development in the autoimmune-prone NOD mouse model was directly impaired through the use of the CRISPR/Cas9 gene editing system to selectively delete the CNS1 region of *Foxp3*. The results suggest that polyclonal tTregs are sufficient to control T1D. In parallel, another group generated a NOD CNS1^−/−^ model using the CRISPR/Cas9 system. Their results differed, suggesting that pTregs were indeed critical for control of autoimmune NOD diabetes, with NOD CNS1^−/−^ males and females exhibiting higher rates of T1D incidence [47]. Our results suggest that tTregs are the dominant regulators of T1D in the NOD mouse model, but under certain conditions such as targeted antigen-specificities or certain external factors, pTregs may play a role in the progression of T1D.

## Results

### CNS1 deletion in NOD mice inhibits *in vitro* iTreg generation

The CNS1 region of *Foxp3* was directly deleted in the NOD mouse background using the CRISPR-Cas9 genome editing system (Fig 1A). Single guide RNAs (sgRNAs) were generated based on the 705 bp region that spans the regulatory element previously described in the B6 *Foxp3* locus, namely from +2003 to +2707 bp from the transcriptional start site of *Foxp3* [38]. PCR and DNA sequencing confirmed the deletion of a 736 bp sequence (+1976 to +2711), which spans the CNS1 region (Fig 1B, S1 Fig). To verify that the genetic deletion corresponded with a functional defect, we performed an *in vitro* Foxp3 induction assay using TGF-β. This assay demonstrated an impairment in the generation of iTregs (Fig 1C), confirming that genetic deletion of the CNS1 region in the NOD background conferred the previously described functional defect observed in the B6 background.

**Fig 1.**
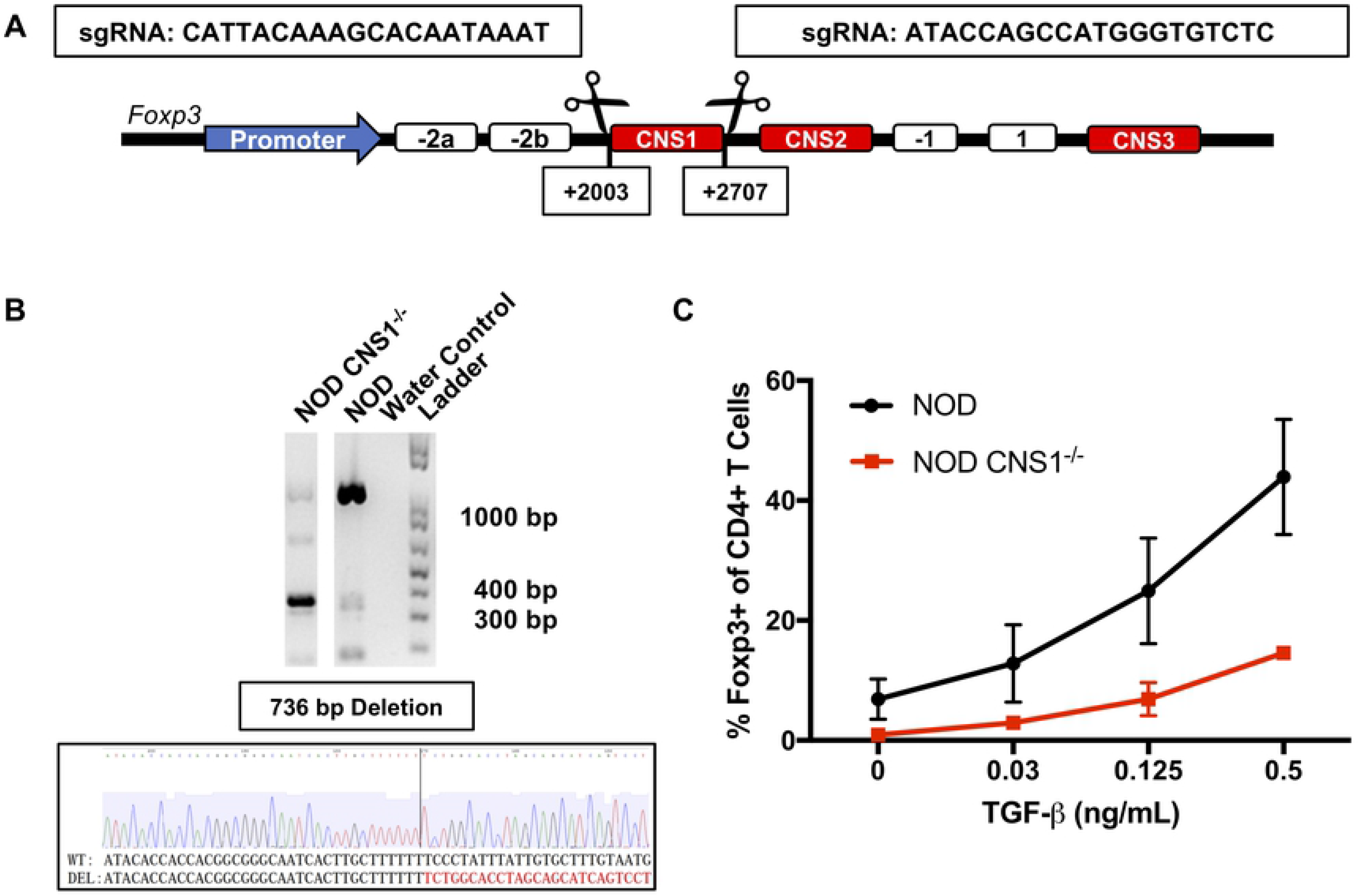
Design, generation, and verification of CNS1^−/−^ in the NOD mouse background. (A) Schematic representation of the Foxp3 locus highlighting the originally defined CNS1 region (+2003 - +2707 from transcription start site) and corresponding sgRNA sequences for CRISPR/Cas9 targeted cutting. (B) Genotyping PCR showing deletion of the CNS1 region in the primary founder and genetic sequencing after CRISPR/Cas9 deletion verifying a 736 base pair deletion in the Foxp3 locus. (C) CD4^+^CD8^−^CD25^−^CD62L^hi^ naïve T cells were sorted from NOD and NOD CNS1^−/−^ mice and stimulated in 96-well plates with plate-coated anti-CD3 (5 μg/mL) and anti-CD28 (0.5 μg/mL) antibodies for 72 hours with hIL-2 (100 IU/mL) in the absence or presence of TGF-β. Cells were than stained for CD4, CD25, and Foxp3 and analyzed with flow cytometry (n=2 mice per genotype, two technical replicates per mouse used).

### Treg quantitation and phenotyping in NOD CNS1^−/−^ mice

In order to directly address the specific consequences of *Foxp3* CNS1 deletion on pTreg generation, pre-diabetic female NOD mice aged 8-13 weeks were sacrificed and the *in vivo* Treg compartment was analyzed. Overall Treg percentages were comparable between NOD and NOD CNS1^−/−^ mice in both lymphoid and non-lymphoid tissues, including the thymus, spleen, mesenteric lymph nodes (mLN), pancreatic islets, and large intestine lamina propria (LI LP), with a small but significant decrease in Treg percentages in the pancreatic lymph nodes of NOD CNS1^−/−^ females (Fig 2A).

**Fig 2.**
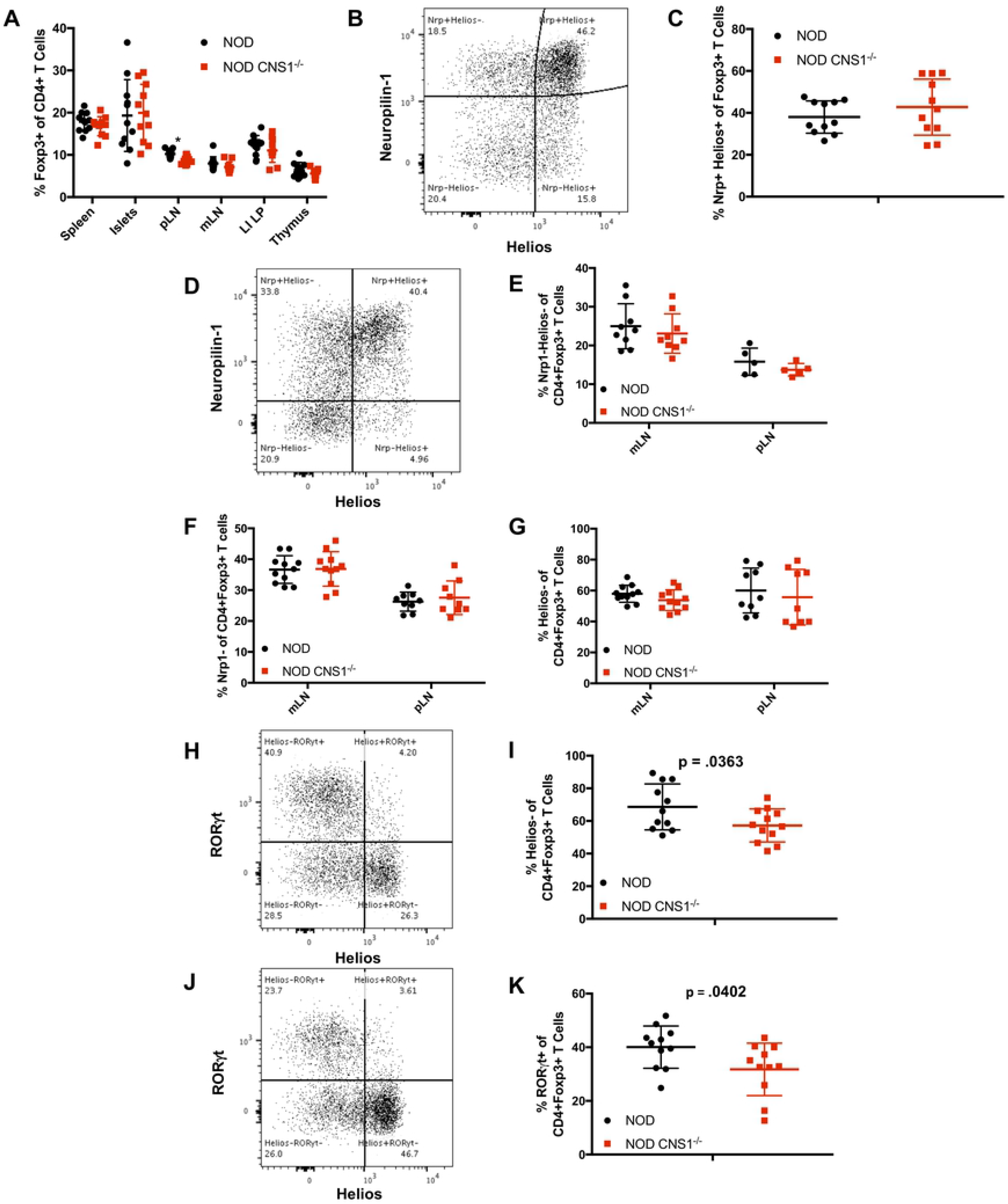
Broad characterization of lymphoid and non-lymphoid tissues in NOD and NOD CNS1^−/−^ mice. (A) Quantification of Treg percentages in various tissues (pLN = pancreatic lymph node, mLN = mesenteric lymph node, LI LP = large intestine lamina propria). (B) Representative plot of CD4+CD8-Foxp3+ thymocytes showing Neuropilin-1 (Nrp11) and Helios expression. (C) Quantification of Nrp1+Helios+ CD4+CD8-Foxp3+ Treg thymocytes. (D) Representative plot of CD4+Foxp3+ T cells expressing Nrp1 and Helios in the mesenteric lymph node (mLN). Quantification of Nrp1-Helios- (E), Nrp1- (F), and Helios- (G) Tregs in mLN and pancreatic lymph node (pLN), reflecting potential pTreg populations. Representative plot of CD4+Foxp3+ T cells expressing RORγt and Helios in the large intestine lamina propria (LI LP) of NOD (H) and NOD CNS1^−/−^ (I) mice. Quantification of Helios- (J) and RORγt+ (K) Tregs in the LI LP, reflecting potential pTreg populations. All figures show NOD and NOD CNS1^−/−^ female mice, 8-13 weeks, n=11.

We also characterized the Tregs present based on markers such as Nrp1 and Helios expression. As expected, Nrp1 and Helios expression within the thymus were comparable between NOD and NOD CNS1^−/−^ females (Fig 2B-C). We examined whether a decrease in the Nrp1^−^ and Helios^−^ populations in the periphery would be evident based on previous results in B6 mice (Fig 2D). However, there was no significant difference in the percentage of Nrp1^−^Helios^−^ (Fig 2E), Nrp1^−^ (Fig 2F), or Helios^−^ (Fig 2G) Tregs in the mLN and pLN.

Previous studies have suggested that the LI LP is largely comprised of Helios^−^ pTregs, with a majority being positive for the transcription factor retinoic acid-related orphan receptor-γt (RORγt) [48,49]. These RORγt+ Tregs are induced by the presence of complex microbiota and provide anti-inflammatory functions [50] and have been shown to have a distinct repertoire [51]. For example, RORγt^+^ pTregs responsive to the short-chain fatty acid (SCFA) butyrate are generated in the gut [42] where they function to establish border-dwelling bacteria that protect against microbial exposure and normal metabolite profiles [43]. Therefore, we hypothesized that there would be a reduction in the Helios^−^ and RORγt^+^ pTreg populations in NOD CNS1^−/−^ females. Cells expressing Helios versus RORγt were mutually exclusive in Tregs isolated from the gut of both NOD (Fig 2H) and NOD CNS1^−/−^ (Fig 2J) mice. There was a small but significant shift in the percentage of the Helios^−^ as well as RORγt^+^ Treg populations in NOD CNS1^−/−^ mice (Fig 2I and Fig 2K, respectively). The implications of these differences are unclear and may reflect changes in the ability of the pTregs to develop in response to exposure to microbiota.

### Islet antigen-reactive Tconv and Treg cells are unchanged in NOD CNS1^−/−^ mice

Previous studies have shown that defects in Tregs in local tissues result in the expansion of autoreactive Tconv cells. Therefore, we examined the relative number of antigen-specific T cells in the pancreatic islets of NOD and NOD CNS1^−/−^ female mice. To assess antigen-specific T cells, I-A^g7^-peptide tetramers was used to stain cells specific for two insulin beta chain (InsB:9-23) variants: p8E, which represents the natural InsB peptide sequence; and p8G, which contains a mutation mimicking a proposed C-terminal InsB:9-20 truncation that shifts the MHC register of presentation for the epitope [52]. Insulin peptide is thought to be produced within the thymus by medullary thymic epithelial cells (mTECs) expressing the autoimmune regulator (Aire) transcription factor, driving thymic selection of T cells [53]. I-A^g7^-peptide tetramers were also used to stain for reactivity against post-transcriptional HIPs generated from the fusion products of proinsulin C-peptide with chromogranin A (ChgA) or islet amyloid polypeptide 2 (IAPP2) [25].

MHC II-islet peptide-specific tetramer staining revealed that Tconvs were more reactive to the HIP peptides than InsB peptides (Fig 3A). This result is consistent with the idea that InsB-reactive Tconvs would be deleted within the thymus while the so-called HIPs could represent peripherally-derived neoantigens that escape negative selection. Moreover, when comparing tetramer staining in Tconvs between NOD and NOD CNS1^−/−^ females, no statistical difference was observed between the percentage of cells reactive to either the InsB:9-23 p8E (Fig 3C) or p8G variants (Fig 3D). The same was true for the Ins:ChgA 2.5HIP (Fig 3E) and the Ins:IAPP2 6.9HIP (Fig 3F) tetramers.

**Fig 3.**
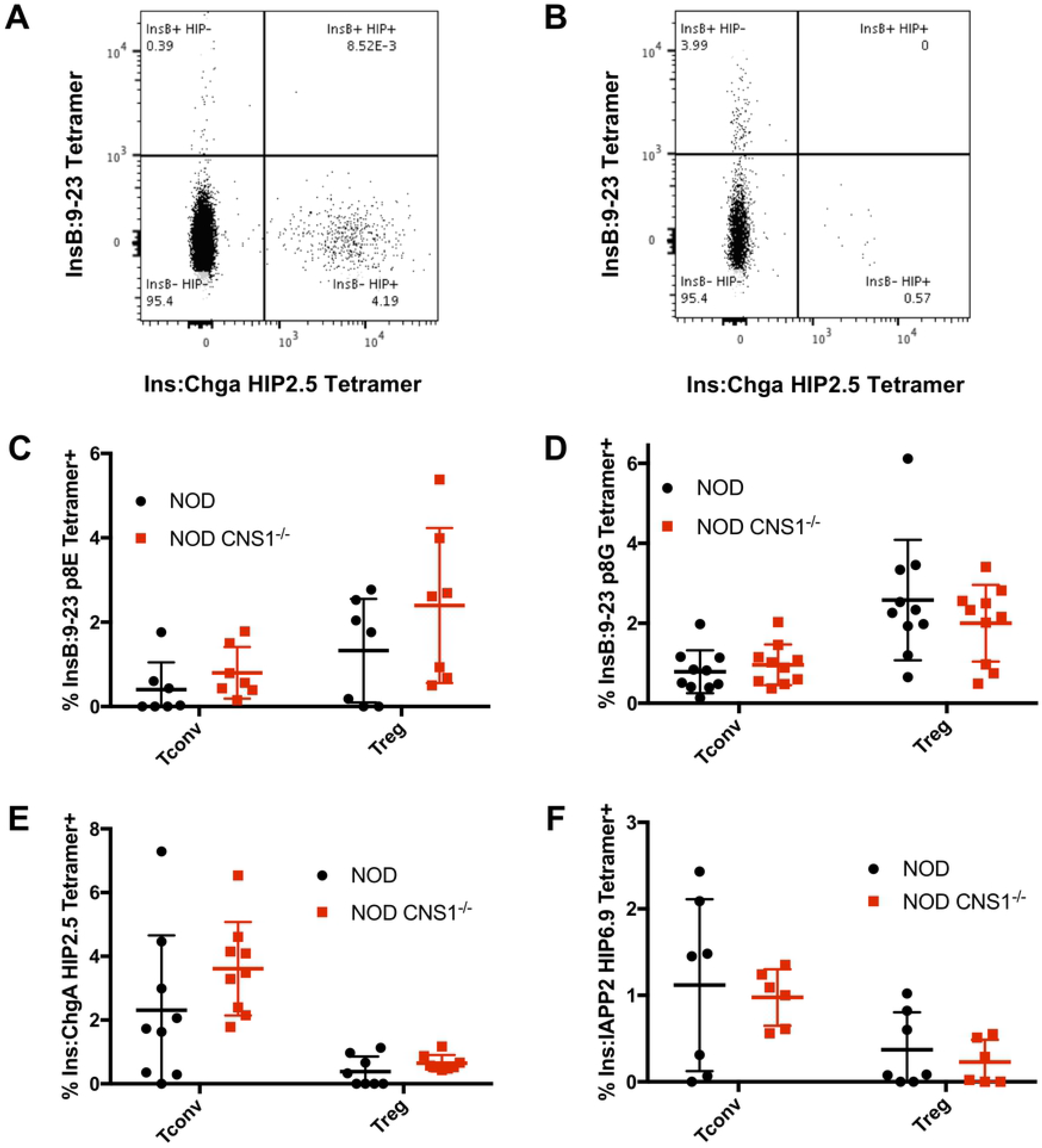
Autoreactive Treg and Tconv percentages between NOD and NOD CNS1^−/−^ mice. Representative tetramer staining plots of CD4+Foxp3− Tconvs (A) and CD4+Foxp3+ Tregs (B) for insulin peptide InsB 9:23 and hybrid insulin peptide (HIP) fusion of insulin and chromogranin A show differential tetramer binding between Tregs and Tconvs. Tetramer staining within pancreatic islets for InsB:9-23 p8E (n=7) (C) and InsB:9-20 p8G (n=10) (D) variants and Ins:ChgA HIP2.5 (n=9) (E) and Ins:IAPP2 HIP6.9 (n=7) (F) is quantified and compared between NOD and NOD CNS1^−/−^ pre-diabetic females aged 8-13 weeks (* = p < .05; ** = p < .005).

Thus, we hypothesized that HIP-reactive Tregs would represent a pTreg population derived from HIP-reactive Tconvs, which would predict that there would be fewer HIP-reactive Tregs in the islets of NOD CNS1^−/−^ mice. However, Tregs isolated from pancreatic islets harvested from wild type NOD mice did not react with either the HIP2.5 or the HIP6.9 tetramers, suggesting that there may not be a significant percentage of pTregs in the islets overall. In contrast, Tregs reactive with the p8E and p8G tetramers were higher as compared to Tconvs (Fig 3B). Furthermore, there was no statistical difference observed between NOD and NOD CNS1^−/−^ females in the percentage of Tregs reactive to the p8E (Fig 3C), p8G (Fig 3D), 2.5HIP (Fig 3E), and 6.9HIP (Fig 3F) tetramers. These data support the idea that CNS1 deletion does not alter the relative percentage of autoreactive Tconvs or Tregs within the pancreatic islets. Furthermore, the Treg compartment within the pancreatic islets seem to be comprised of largely tTregs.

### No difference in glucose tolerance, time of T1D onset, or disease incidence in NOD CNS1^−/−^ mice despite differences in insulitis

The ultimate potential consequence of the effects of disruption of pTreg induction was read out by the impact on T1D development in the NOD background. Female NOD and NOD CNS1^−/−^ pancreata were collected at 6, 10, and 14 weeks of age for histology to measure insulitis (Fig 4A). Histological analysis of the isolated islets showed that there was some increase in insulitis in NOD CNS1^−/−^ mice as early as 6 weeks and 10 weeks of age in the CNS1^−/−^ mice. However, by 14 weeks, the time of initial development of clinical diabetes in our colony, there was only a marginal difference in insulitis between NOD and NOD CNS1^−/−^ females, suggesting that the deletion of CNS1 did not ultimately lead to an increase in beta cell destruction. In fact, in younger pre-diabetic NOD CNS1^−/−^ female mice, normal glucose tolerance was maintained (Fic 4B-C). These results show that the small differences in insulitis between NOD and NOD CNS1^−/−^ mice at earlier ages were not consequential for either acute or chronic changes in blood glucose control. Importantly, these early differences in insulitis did not influence the onset or incidence of T1D in either female or male NOD CNS1^−/−^ mice, which was not statistically different between the groups (Fig 4D).

**Fig 4.**
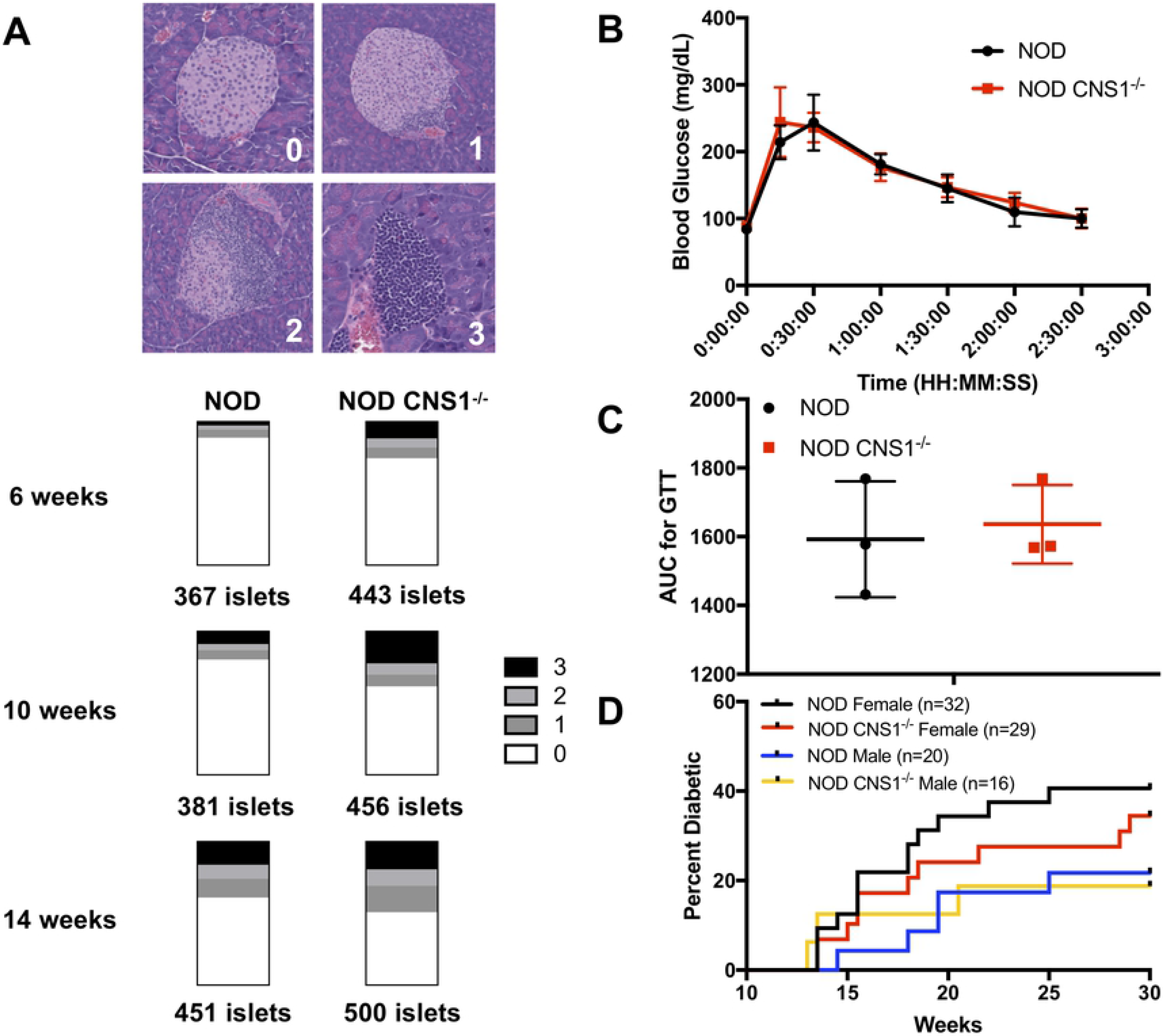
Comparison of T1D development and glucose tolerance between NOD and NOD CNS1^−/−^ mice. (A) Pooled insulitis scoring of female NOD and NOD CNS1^−/−^ mice at 6, 10, and 14 weeks of age (n=3 mice per age and genotype). Representative H&E histology images of pancreatic islets illustrating insulitis scoring rubric are shown at the top (0 = no insulitis, 1 = peri-insulitis or <25% infiltration, 2 is 25%-75% infiltration, and 3 is >75% infiltration or full islet destruction). p < 0.0001 for 6 weeks and 10 weeks, and p = 0.0140 at 14 weeks based on a chi-square comparison of islets scored as 0 to islets scored as 1, 2, or 3. (B) Glucose tolerance testing of 8-9 week old NOD and NOD CNS1^−/−^ females after 16 hour overnight fast (n=3 per group) (C) Glucose area under the curve (AUC) of glucose tolerance testing shown in (B). (D) Diabetes incidence based on weekly blood glucose measurements comparing NOD and NOD CNS1^−/−^ males and females.

## Discussion

Our data show that deletion of CNS1 in the NOD background does not impact Treg frequencies or expression of known markers associated with pTregs and tTregs. Furthermore, deletion does not change the frequency of islet antigen-specific Tregs or Tconvs within the pancreatic islets. This includes InsB:9-23, a self-antigen known to be expressed in the thymus, and HIP tetramers, specific for post-translationally fused peptides generated in the periphery. These results are consistent with recent reports that show that InsB:9-23 tetramer binding is enriched in Tregs while HIP tetramer binding is enriched in Tconvs [23,26]. Finally, there is no difference in glucose tolerance, diabetes onset or incidence when comparing the NOD and NOD CNS1^−/−^ mice.

These results are consistent with another group who generated a Smad3 binding mutation within CNS1 of NOD mice, which led to an age-dependent decrease in Treg frequency within the LI LP and other gut associated lymphoid tissues (GALT) but did not affect susceptibility to colitis [45]. However, another group independently generated a NOD CNS1^−/−^ model using different sgRNAs [47]. Their work revealed a similar cellular phenotype where the frequency of Tregs found in most lymphoid and non-lymphoid tissues remained largely the same, but they observed a small but significant decrease in the Treg frequencies within the LI LP instead of the pLN. Interestingly, this alternate NOD CNS1^−/−^ model, there were significant reductions in Helios expression in Tregs in more lymphoid and non-lymphoid tissues than what we observed. Moreover, the alternate NOD CNS1^−/−^ model showed a much starker difference in insulitis at 6 and 10 weeks of age, and although there is not a significant difference in insulitis by 14 weeks of age, overall insulitis was much higher in both their NOD and NOD CNS1^−/−^ females. These results suggest that there may be a role for pTregs early in disease progression but overtime, the tTregs dominate and control disease progression such that depending on the colony and the degree of early beta cell damage, the resulting ability to control disease and maintain normoglycemia may be different. In the case of our mice, tTregs control disease progression preventing clinical disease, while in the other NOD *Foxp3* CNS1 deletion model a clinical disease phenotype is evident in both male and female mice.

Although many of the results we obtained are consistent with the previous NOD CNS1^−/−^ studies, there are a number of factors that could account for both earlier and more severe insulitis in the CNS1^−/−^ mice described by Kissler and colleagues than those in our colony. One difference between the different NOD CNS1^−/−^ mouse strains is that the CNS1 deletion is not identical. Kissler and colleagues used sgRNAs that span a wider region around the DNA than the ones we utilized. Thus, the resulting edit generated a slightly larger 795 bp deletion (+1949 to +2743 from the *Foxp3* transcriptional start site) (unpublished information from Kissler) than was generated in the animals herein (+1976 to +2711). Whether these additional nucleotides might alter DNA binding of other enhancer proteins remains unclear but might be impactful.

Perhaps the most likely difference between the two mouse models resides in the gut microbiome or other environmental factors. There is extensive literature documenting that innate microbial sensing and gut microbial constituents can shift T1D incidence, explaining differences observed in diabetes incidence among animal facilities [54,55]. Specific differences in the gut microbial community of NOD males and females post-puberty also contribute to the differences in T1D incidence seen between the sexes [56,57], and microbiome composition can be used to predict development of T1D [58]. The gut microbiome might be linked to T1D due to the fact that the pLN is also a draining site for microbial antigens [59], and autoreactive cells have also shown cross-reactivity between microbial antigens and islet antigens [60,61]. In fact, recent characterization of the B6 CNS1^−/−^ mouse directly revealed that defects in gut pTreg generation alter gut microbiome colonization [42,43]. Since there is a change in gut Tregs in both NOD CNS1^−/−^ models generated, the ability to observe a similar disease phenotype in our NOD CNS1^−/−^ model might be masked by differences in gut microbiome composition. Future studies that sequence and compare the microbiome of NOD and NOD CNS1^−/−^ mice generated in the two facilities would be required to address this issue directly.

In summary, our data suggest that tTregs play the dominant role in the control of islet-specific autoimmunity and subsequent development of diabetes. This conclusion is supported by the observation that within the pancreatic islets, there are not many Tregs that recognize tissue-specific, peripherally-generated autoantigens as detected by HIP I-A^g7^ tetramers, consistent with the notion that the Treg compartment within the islets is largely comprised of tTregs. Although our work shows that tTregs are sufficient for controlling T1D in NOD mice, the role of CNS1-dependent pTregs might be influenced by environmental factors based on the differences seen between our model and an independently-generated NOD CNS1 deletion. This observation is consistent with what is known about T1D and could provide further insight into the interplay between the immune system and environment that determines both mouse and human disease susceptibility and outcomes.

## Methods

### Mice

NOD CNS1^−/−^ mice were made by the JAX Model Generation services using the CRISPR/Cas9 system (gRNAs 5’-CATTACAAAGCACAATAAAT-3’ and 5’-ATACCAGCCATGGGTGTCTC-3’). Potential founders were genotyped and sequenced using the following primers: 5’-CAGCAGTGCTCTTACCCATG-3’ and 5’-CAGTGAGAGCAGTTTAGAGG-3’. Founder mouse progeny were crossed to NOD JAX mice for at least three generations before experiments to account for potential off-target CRISPR/Cas9 effects. Hyperglycemia and spontaneous induction of T1D in NOD mice were monitored by weekly blood glucose measurements made with the OneTouch Ultra 2 glucometer and UNISTRIP1 generic blood glucose test strips. Mice with two consecutive blood glucose readings of at least 250 mg/dL were considered diabetic. All experiments were conducted under a protocol approved by the Institutional Animal Care and Use Committee.

### Isolation of lymphocytes and cell staining

Lymph nodes were physically dissociated and filtered to generate single-cell suspensions. Spleens were physically dissociated and incubated at room temperature in ACK red blood cell lysis buffer before filtration. Large intestine lamina propria (LI LP) were digested using the Lamina Propria Dissociation Kit (MACS Miltenyi Biotec) per manufacturer’s instructions. Islet isolation was done courtesy of the UCSF Mouse Islet Isolation Facility [62] and then dissociated in Cell Dissociation Buffer Enzyme-Free PBS-based (ThermoFisher Scientific) at 37°C in a tissue culture incubator. Antibodies used for flow cytometry staining are listed in S1 Table. Cell viability was measured before flow cytometry using the Vi-Cell XR Cell Viability Analyzer (Beckman Coulter) and tracked during flow cytometry using the LIVE/DEAD Fixable Blue Dead Cell Stain Kit for UV Excitation (ThermoFisher Scientific). After viability staining, tetramer staining was performed at room temperature for 1 hour, and then without washing, concentrated surface stain was added for an additional 15 minutes at room temperature. The following mouse I-A^g7^ tetramers were used courtesy of the NIH Tetramer Core Facility: InsB p8E variant (HLVERLYLVCGEEG) conjugated to APC; InsB p8G variant (HLVERLYLVCGGEG) conjugated to APC; Ins:ChgA 2.5HIP (LQTLALWSRMD) conjugated to APC; Ins:IAPP2 6.9HIP (LQTLALNAARD) conjugated to PE; and DPB1*04:01/DPA1*01:03 control human CLIP 87-101 conjugated to APC, BV421, and PE (PVSKMRMATPLLMQA). Fixation and intracellular staining were done using the eBioscience Foxp3/Transcription Factor Staining Buffer Set (ThermoFisher Scientific) per manufacturer’s instructions. Stained single-cell suspensions were analyzed using a Fortessa flow cytometer running FACSDiva (BD Biosciences). FSC 3.0 files were analyzed and presented with FLOWJO Software (flowjo.com).

### *In vitro* Foxp3 induction assay

Naïve T cells (CD4+CD25-CD62Lhi) were isolated and stained as described above and then FACS-Aria sorted into fetal bovine serum (FBS). Cells were resuspended in complete (penicillin/streptomycin, sodium pyruvate, HEPES, NEAA, and β-mercaptoethanol) DMEM plus 10% FBS with human IL-2 (100 IU/mL) and TGF-β (0, 0.03125, 0.125, and 0.5 ng/mL) and cultured in 96-well plates coated with anti-CD3 (5 μg/mL) and anti-CD28 (0.5 μg/mL) for 72 hours at 37°C in a tissue culture incubator. Cells were than stained for CD4, CD25, and Foxp3 (S1 Table) and analyzed using a Fortessa flow cytometer running FACSDiva (BD Biosciences). FSC 3.0 files were analyzed and presented with FLOWJO Software (flowjo.com).

### Histology and pancreatic insulitis scoring

Pancreata were fixed in 10% neutral buffered formalin solution overnight and dehydrated in 70% ethanol. Histology was performed by HistoWiz Inc. (histowiz.com) using a Standard Operating Procedure and fully automated workflow. Samples were processed, embedded in paraffin, and sectioned at 6 μm with 175 μm steps between slices. Three sections per mouse pancreas were collected. Haemotoxylin and Eosin (H&E) staining was performed, and then sections were dehydrated and film coverslipped using a TissueTek-Prisma and Coverslipper (Sakura). Whole slide scanning (40x) was performed on an Aperio AT2 (Leica Biosystems). Islets were scored blinded to genotype and age with at least 100 islets scored per mouse. Insulitis was scored as none (0), peri-insulitis or less than 25% infiltration (1), 25-75% infiltration (2), or greater than 75% infiltration (3). Islet scoring was done in the open source software QuPath (qupath.github.io) [63].

### Glucose tolerance test

Glucose tolerance was tested in 8-9-week-old NOD and NOD CNS1^−/−^ mice after a 16 hour overnight fast. Mice were challenged with injections of 1M glucose in PBS (2 g glucose/kg of total body weight) intraperitoneally and blood glucose was measured with the OneTouch Ultra 2 glucometer and UNISTRIP1 generic blood glucose test strips at baseline and then 15, 30, 60, 90, 120, and 150 minutes post-challenge.

### Statistics and methods of analysis

Data analyses were performed using GraphPad Prism 7 software (graphpad.com/scientific-software/prism), and values at p < .05 were deemed significant. For comparison of the insulitis scores, the chi-square test was used to calculate if there was a significant difference in the proportion of islets without any peri-insulitis or insulitis (score = 0) to islets with peri-insulitis or full insulitis (score = 1, 2, or 3). All other statistical comparisons were done using a two-tailed, unpaired t test.

## Acknowledgements

The authors thank Michael Lee, Vinh Nguyen, and the UCSF Flow Core for assisting with flow cytometry; Vinh Nguyen and Vi Dang of the UCSF Islet Production Core for providing mouse pancreatic islets; Shen Dong, Arabella Young, and Wendy Rosenthal for conceptual and technical advice for experiments; and Yanli Wang for animal care. D.R.H., F.V.G. and J.A.B. designed the study; D.R.H. performed experiments; and D.R.H. and J.A.B. wrote the manuscript with input from F.V.G.

## Supporting Information

**S1 Fig. Sequence of CNS1 deletion in NOD mice**. Displayed here is the +1960 to +2722 region from the transcriptional start side of Foxp3. The original CNS1 region identified in B6 mice (+2003 to +2707) is highlighted in yellow, while the sequence of the deleted region generated in the NOD background with CRISPR/Cas9 (+1976 to +2711) is bolded in red.

**S2 Table. Reference list of flow cytometry antibodies used for analysis.**

## References List

1. Sakaguchi, Shimon; Sakaguchi, Noriko; Asano Masanao; Itoh, Misako; Toda M. Immunologic Self-Tolerance Maintained by activated T cells expressing IL-2 Receptor a-chains (CD25). J Immunol. 1995;

2. Takahashi T, Tagami T, Yamazaki S, Uede T, Shimizu J, Sakaguchi N, et al. Immunologic Self-Tolerance Maintained by Cd25 ^+^ Cd4 ^+^ Regulatory T Cells Constitutively Expressing Cytotoxic T Lymphocyte–Associated Antigen 4. J Exp Med. 2000;

3. Hori S, Nomura T, Sakaguchi S. Control of regulatory T cell development by the transcription factor Foxp3.[see comment]. Science. 2003;299(5609):1057–61.

4. Fontenot JD, Gavin MA, Rudensky AY. Foxp3 programs the development and function of CD4^+^ CD25^+^ regulatory T cells. Nat Immunol. 2003;

5. Fontenot JD, Rasmussen JP, Williams LM, Dooley JL, Farr AG, Rudensky AY. Regulatory T cell lineage specification by the forkhead transcription factor Foxp3. Immunity. 2005;22(3):329–41.

6. Salomon B, Lenschow DJ, Rhee L, Ashourian N, Singh B, Sharpe A, et al. B7/CD28 costimulation is essential for the homeostasis of the CD4+CD25+immunoregulatory T cells that control autoimmune diabetes. Immunity. 2000;12(4):431–40.

7. Wildin RS, Ramsdell F, Peake J, Faravelli F, Casanova JL, Buist N, et al. X-linked neonatal diabetes mellitus, enteropathy and endocrinopathy syndrome is the human equivalent of mouse scurfy. Nat Genet. 2001;27(1):18–20.

8. Khattri R, Cox T, Yasayko SA, Ramsdell F. An essential role for Scurfin in CD4+CD25+T regulatory cells. J Immunol. 2017;198(3):993–8.

9. Brode S, Raine T, Zaccone P, Cooke A. Cyclophosphamide-Induced Type-1 Diabetes in the NOD Mouse Is Associated with a Reduction of CD4+CD25+Foxp3+ Regulatory T Cells. J Immunol [Internet]. 2006;177(10):6603–12. Available from: http://www.jimmunol.org/cgi/doi/10.4049/jimmunol.177.10.6603

10. Feuerer M, Shen Y, Littman DR, Benoist C, Mathis D. How Punctual Ablation of Regulatory T Cells Unleashes an Autoimmune Lesion within the Pancreatic Islets. Immunity [Internet]. 2009;31(4):654–64. Available from: http://dx.doi.org/10.1016/j.immuni.2009.08.023

11. Jordan MS, Boesteanu A, Reed AJ, Petrone AL, Holenbeck AE, Lerman MA, et al. Thymic selection of CD4+CD25+ regulatory T cells induced by an agonist self-peptide. Nat Immunol. 2001;2(4):301–6.

12. Apostolou I, Sarukhan A, Klein L, Boehmer H Von. Origin of regulatory T cells with known specificity for antigen. 2002;3(8):2–9.

13. Chen W, Jin W, Hardegen N, Lei K, Li L, Marinos N, et al. Conversion of Peripheral CD4 + CD25 − Naive T Cells to CD4 + CD25 + Regulatory T Cells by TGF-β Induction of Transcription Factor Foxp3. J Exp Med. 2003;198(12):1875–86.

14. Benson MJ, Pino-Lagos K, Rosemblatt M, Noelle RJ. All-trans retinoic acid mediates enhanced T reg cell growth, differentiation, and gut homing in the face of high levels of co-stimulation. J Exp Med. 2007;204(8):1765–74.

15. Coombes JL, Siddiqui KRR, Arancibia-Cárcamo C V., Hall J, Sun C-M, Belkaid Y, et al. A functionally specialized population of mucosal CD103 ^+^ DCs induces Foxp3 ^+^ regulatory T cells via a TGF-β– and retinoic acid–dependent mechanism. J Exp Med [Internet]. 2007;204(8):1757–64. Available from: http://www.jem.org/lookup/doi/10.1084/jem.20070590

16. Sun C-M, Hall JA, Blank RB, Bouladoux N, Oukka M, Mora JR, et al. Small intestine lamina propria dendritic cells promote de novo generation of Foxp3 T reg cells via retinoic acid. J Exp Med [Internet]. 2007;204(8):1775–85. Available from: http://www.jem.org/lookup/doi/10.1084/jem.20070602

17. Apostolou I, von Boehmer H. In Vivo Instruction of Suppressor Commitment in Naive T Cells. J Exp Med [Internet]. 2004;199(10):1401–8. Available from: http://www.jem.org/lookup/doi/10.1084/jem.20040249

18. Mucida D, Kutchukhidze N, Erazo A, Russo M, Lafaille JJ, Curotto De Lafaille MA. Oral tolerance in the absence of naturally occurring Tregs. J Clin Invest. 2005;

19. Kretschmer K, Apostolou I, Hawiger D, Khazaie K, Nussenzweig MC, von Boehmer H. Inducing and expanding regulatory T cell populations by foreign antigen. Nat Immunol. 2005;6(12):1219–27.

20. Feuerer M, Jiang W, Holler PD, Satpathy A, Campbell C, Bogue M, et al. Enhanced thymic selection of FoxP3 %. regulatory T cells in the NOD mouse model of autoimmune diabetes. 2007;104(46):18181–6.

21. Ferreira C, Palmer D, Blake K, Garden OA, Dyson J. Reduced Regulatory T Cell Diversity in NOD Mice Is Linked to Early Events in the Thymus. J Immunol. 2014;

22. Kern J, Drutel R, Leanhart S, Bogacz M, Pacholczyk R. Reducti on of T cell receptor diversity in NOD mice prevents development of type 1 diabetes but not Sjögren’s syndrome. PLoS One. 2014;9(11).

23. Spence A, Purtha W, Tam J, Dong S, Kim Y, Ju C, et al. Revealing the specificity of regulatory T cells in murine autoimmune diabetes. Proc Natl Acad Sci [Internet]. 2018;115(24):201808331. Available from: http://www.pnas.org/lookup/doi/10.1073/pnas.1808331115

24. Delong T, Baker RL, He J, Barbour G, Bradley B, Haskins K. Diabetogenic T-cell clones recognize an altered peptide of chromogranin A. Diabetes. 2012;61(12):3239–46.

25. Delong T, Wiles TA, Baker RL, Bradley B, Barbour G, Reisdorph R, et al. Pathogenic CD4 T cells in type 1 diabetes recognize epitopes formed by peptide fusion. Science (80-). 2016;351(6274):711–4.

26. Baker RL, Jamison BL, Wiles TA, Lindsay RS, Barbour G, Bradley B, et al. CD4 T cells reactive to hybrid insulin peptides are indicators of disease activity in the NOD mouse. Diabetes. 2018;67(9):1836–46.

27. Mannering SI, Harrison LC, Williamson NA, Morris JS, Thearle DJ, Jensen KP, et al. The insulin A-chain epitope recognized by human T cells is posttranslationally modified. J Exp Med [Internet]. 2005;202(9):1191–7. Available from: http://www.jem.org/lookup/doi/10.1084/jem.20051251

28. Van Lummel M, Duinkerken G, Van Veelen PA, De Ru A, Cordfunke R, Zaldumbide A, et al. Posttranslational modification of HLA-DQ binding islet autoantigens in type 1 diabetes. Diabetes. 2014;63(1):237–47.

29. McGinty JW, Chow IT, Greenbaum C, Odegard J, Kwok WW, James EA. Recognition of posttranslationally modified GAD65 epitopes in subjects with type 1 diabetes. Diabetes. 2014;63(9):3033–40.

30. Haribhai D, Williams JB, Jia S, Nickerson D, Schmitt EG, Edwards B, et al. A requisite role for induced regulatory T cells in tolerance based on expanding antigen receptor diversity. Immunity [Internet]. 2011;35(1):109–22. Available from: http://dx.doi.org/10.1016/j.immuni.2011.03.029

31. Yadav M, Louvet C, Davini D, Gardner JM, Martinez-Llordella M, Bailey-Bucktrout S, et al. Neuropilin-1 distinguishes natural and inducible regulatory T cells among regulatory T cell subsets in vivo. J Exp Med [Internet]. 2012;209(10):1713–22. Available from: http://www.jem.org/lookup/doi/10.1084/jem.20120822

32. Weiss JM, Bilate AM, Gobert M, Ding Y, Curotto de Lafaille MA, Parkhurst CN, et al. Neuropilin 1 is expressed on thymus-derived natural regulatory T cells, but not mucosa-generated induced Foxp3 ^+^ T reg cells. J Exp Med [Internet]. 2012;209(10):1723–42. Available from: http://www.jem.org/lookup/doi/10.1084/jem.20120914

33. Thornton AM, Korty PE, Tran DQ, Wohlfert EA, Murray PE, Belkaid Y, et al. Expression of Helios, an Ikaros Transcription Factor Family Member, Differentiates Thymic-Derived from Peripherally Induced Foxp3 + T Regulatory Cells. J Immunol. 2010;184(7):3433–41.

34. Akimova T, Beier UH, Wang L, Levine MH, Hancock WW. Helios expression is a marker of T cell activation and proliferation. PLoS One. 2011;6(8).

35. Gottschalk RA, Corse E, Allison JP. Expression of Helios in Peripherally Induced Foxp3+ Regulatory T Cells. J Immunol [Internet]. 2012;189(2):500–500. Available from: http://www.jimmunol.org/cgi/doi/10.4049/jimmunol.1290033

36. Szurek E, Cebula A, Wojciech L, Pietrzak M, Rempala G, Kisielow P, et al. Differences in expression level of Helios and neuropilin-1 do not distinguish thymus-derived from extrathymically-induced CD4+Foxp3+ regulatory T cells. PLoS One. 2015;10(10):1–16.

37. Nutsch K, Chai JN, Ai TL, Russler-Germain E, Feehley T, Nagler CR, et al. Rapid and Efficient Generation of Regulatory T Cells to Commensal Antigens in the Periphery. Cell Rep [Internet]. 2016;17(1):206–20. Available from: http://dx.doi.org/10.1016/j.celrep.2016.08.092

38. Zheng Y, Josefowicz S, Chaudhry A, Peng XP, Forbush K, Rudensky AY. Role of conserved non-coding DNA elements in the Foxp3 gene in regulatory T-cell fate. Nature [Internet]. 2010;463(7282):808–12. Available from: http://dx.doi.org/10.1038/nature08750

39. Travis MA, Reizis B, Melton AC, Masteller E, Tang Q, Proctor JM, et al. Loss of integrin a v b 8 on dendritic cells causes autoimmunity and colitis in mice. 2007;449(September):361–6.

40. Schallenberg S, Tsai P-Y, Riewaldt J, Kretschmer K. Identification of an immediate Foxp3 ^−^ precursor to Foxp3 ^+^ regulatory T cells in peripheral lymphoid organs of nonmanipulated mice. J Exp Med. 2010;

41. Josefowicz SZ, Niec RE, Kim HY, Treuting P, Chinen T, Zheng Y, et al. Extrathymically generated regulatory T cells control mucosal T H 2 inflammation. Nature [Internet]. 2012;482(7385):395–9. Available from: http://dx.doi.org/10.1038/nature10772

42. Arpaia N, Campbell C, Fan X, Dikiy S, Van Der Veeken J, Deroos P, et al. Metabolites produced by commensal bacteria promote peripheral regulatory T-cell generation. Nature [Internet]. 2013;504(7480):451–5. Available from: http://dx.doi.org/10.1038/nature12726

43. Campbell C, Dikiy S, Bhattarai SK, Chinen T, Matheis F, Calafiore M, et al. Extrathymically Generated Regulatory T Cells Establish a Niche for Intestinal Border-Dwelling Bacteria and Affect Physiologic Metabolite Balance. Immunity [Internet]. 2018;1–13. Available from: https://doi.org/10.1016/j.immuni.2018.04.013

44. Samstein RM, Josefowicz SZ, Arvey A, Treuting PM, Rudensky AY. Extrathymic generation of regulatory T cells in placental mammals mitigates maternal-fetal conflict. Cell [Internet]. 2012;150(1):29–38. Available from: http://dx.doi.org/10.1016/j.cell.2012.05.031

45. Schlenner SM, Weigmann B, Ruan Q, Chen Y, von Boehmer H. Smad3 binding to the foxp3 enhancer is dispensable for the development of regulatory T cells with the exception of the gut. J Exp Med [Internet]. 2012;209(9):1529–35. Available from: http://www.jem.org/lookup/doi/10.1084/jem.20112646

46. Petzold C, Steinbronn N, Gereke M, Strasser RH, Sparwasser T, Bruder D, et al. Fluorochrome-based definition of naturally occurring Foxp3+regulatory T cells of intra- and extrathymic origin. Eur J Immunol. 2014;44(12):3632–45.

47. Schuster C, Jonas F, Zhao F, Kissler S. Peripherally-induced regulatory T cells contribute to the control of autoimmune diabetes in the NOD mouse model. Eur J Immunol [Internet]. 2018;1–6. Available from: http://doi.wiley.com/10.1002/eji.201847498

48. Ohnmacht C, Park J-H, Cording S, Wing JB, Atarashi K, Obata Y, et al. The microbiota regulates type 2 immunity through ROR t+ T cells. Science (80-) [Internet]. 2015;349(6251):989–93. Available from: http://www.sciencemag.org/cgi/doi/10.1126/science.aac4263

49. Sefik E, Geva-Zatorsky N, Oh S, Konnikova L, Zemmour D, McGuire AM, et al. Individual intestinal symbionts induce a distinct population of ROR + regulatory T cells. Science (80-) [Internet]. 2015;349(6251):993–7. Available from: http://www.sciencemag.org/cgi/doi/10.1126/science.aaa9420

50. Yang BH, Hagemann S, Mamareli P, Lauer U, Hoffmann U, Beckstette M, et al. Foxp3+T cells expressing RORγt represent a stable regulatory T-cell effector lineage with enhanced suppressive capacity during intestinal inflammation. Mucosal Immunol. 2016;9(2):444–57.

51. Solomon BD, Hsieh C-S. Antigen-Specific Development of Mucosal Foxp3 ^+^ RORγt ^+^ T Cells from Regulatory T Cell Precursors. J Immunol [Internet]. 2016;197(9):3512–9. Available from: http://www.jimmunol.org/lookup/doi/10.4049/jimmunol.1601217

52. Crawford F, Stadinski B, Jin N, Michels A, Nakayama M, Pratt P, et al. Specificity and detection of insulin-reactive CD4+ T cells in type 1 diabetes in the nonobese diabetic (NOD) mouse. Proc Natl Acad Sci [Internet]. 2011;108(40):16729–34. Available from: http://www.pnas.org/cgi/doi/10.1073/pnas.1113954108

53. Anderson MS, Venanzi ES, Klein L, Chen Z, Berzins SP, Turley SJ, et al. Projection of an Immunological Self Shadow Within the Thymus by the Aire Protein. Science (80-) [Internet]. 2002;298(5997):1395–401. Available from: http://science.sciencemag.org/

54. Pozzilli P, Signore A, Williams AJK, Beales PE. NOD mouse colonies around the world-recent facts and figures. Immunol Today. 1993;

55. Wen L, Ley RE, Volchkov PY, Stranges PB, Avanesyan L, Stonebraker AC, et al. Innate immunity and intestinal microbiota in the development of Type 1 diabetes. Nature. 2008;455(7216):1109–13.

56. Markle JGM, Frank DN, Mortin-Toth S, Robertson CE, Feazel LM, Rolle-Kampczyk U, et al. Sex differences in the gut microbiome drive hormone-dependent regulation of autoimmunity. Science. 2013;

57. Yurkovetskiy L, Burrows M, Khan AA, Graham L, Volchkov P, Becker L, et al. Gender bias in autoimmunity is influenced by microbiota. Immunity [Internet]. 2013;39(2):400–12. Available from: http://dx.doi.org/10.1016/j.immuni.2013.08.013

58. Hu Y, Peng J, Li F, Wong FS, Wen L. Evaluation of different mucosal microbiota leads to gut microbiota-based prediction of type 1 diabetes in NOD mice. Sci Rep. 2018;8(1):1–13.

59. Turley SJ, Lee J-W, Dutton-Swain N, Mathis D, Benoist C. Endocrine self and gut non-self intersect in the pancreatic lymph nodes. Proc Natl Acad Sci [Internet]. 2005;102(49):17729–33. Available from: http://www.pnas.org/cgi/doi/10.1073/pnas.0509006102

60. Tai N, Peng J, Liu F, Gulden E, Hu Y, Zhang X, et al. Microbial antigen mimics activate diabetogenic CD8 T cells in NOD mice. J Exp Med [Internet]. 2016;213(10):2129–46. Available from: http://www.jem.org/lookup/doi/10.1084/jem.20160526

61. Culina S, Lalanne AI, Afonso G, Cerosaletti K, Pinto S, Sebastiani G, et al. Islet-reactive CD8 ^+^ T cell frequencies in the pancreas, but not in blood, distinguish type 1 diabetic patients from healthy donors. Sci Immunol [Internet]. 2018;3(20):eaao4013. Available from: http://immunology.sciencemag.org/lookup/doi/10.1126/sciimmunol.aao4013

62. Szot GL, Koudria P, Bluestone JA. Murine Pancreatic Islet Isolation. 2007;1640:7–8.

63. Bankhead P, Loughrey MB, Fernández JA, Dombrowski Y, McArt DG, Dunne PD, et al. QuPath: Open source software for digital pathology image analysis. Sci Rep. 2017;7(1):1–7.

